# Most ventral pallidal cholinergic neurons are cortically projecting bursting basal forebrain cholinergic neurons

**DOI:** 10.1101/2025.02.23.639747

**Authors:** Dániel Schlingloff, Írisz Szabó, Éva Gulyás, Bálint Király, Réka Kispál, Marcus Stephenson-Jones, Balázs Hangya

## Abstract

The ventral pallidum (VP) lies at the intersection of basal ganglia and basal forebrain circuitry, possessing attributes of both major subcortical systems. Basal forebrain cholinergic neurons are rapidly recruited by reinforcement feedback and project to cortical and subcortical forebrain targets; in contrast, striatal cholinergic cells are local interneurons exhibiting classical ‘pause-burst’ responses to rewards. However, VP cholinergic neurons (VPCNs) are less characterized, and it is unclear whether basal forebrain and striatal type cholinergic neurons mix in the VP. Therefore, we performed anterograde and mono-transsynaptic retrograde labeling, in vitro acute slice recordings and bulk calcium recordings of VPCNs. We found that VPCNs broadly interact with the affective circuitry that processes rewards and punishments, targeting the basolateral amygdala, the nucleus accumbens, the medial prefrontal cortex and the lateral habenula, while receiving inputs from the nucleus accumbens, hypothalamus, central amygdala, bed nucleus of stria terminalis and the ventral tegmental area. Bulk calcium recordings revealed VPCN responses to rewards, punishments and reward-predicting cues, like those of the horizontal diagonal band of Broca of the basal forebrain. Acute slice recordings showed that most VPCNs resembled the bursting type of basal forebrain cholinergic neurons (BFCNs), while a few of them were of the regular rhythmic type, which sharply differentiated most VPCNs from striatal cholinergic interneurons. These results were confirmed by in vivo electrophysiological recordings of putative VPCNs largely resembling bursting BFCNs. We conclude that most VPCNs are BFCNs with specialized connectivity to relay aversive and appetitive stimuli to the reinforcement circuitry, possibly implicated in mood disorders and addiction.

## Introduction

The ventral pallidum is considered as the major output structure of the ventral basal ganglia (Maurice et al., 1997; Kupchik et al., 2015; Ahrens et al., 2016; Richard et al., 2016a; Stephenson-Jones et al., 2020), thought to mediate the reinforcing and incentive properties of reward-predicting cues and rewards (Tindell et al., 2005; Tachibana and Hikosaka, 2012; Ahrens et al., 2018; Fujimoto et al., 2019; Ottenheimer et al., 2020a; Hegedüs et al., 2021) and drive reward-seeking behaviors (Smith et al., 2009; Richard et al., 2016a, 2018; Ottenheimer et al., 2020b; Stephenson-Jones et al., 2020; Hernandez-Jaramillo et al., 2024). A recent study further suggested that the VP is monitoring information for upcoming choice behaviors, which it then relays to downstream decision making areas (Ito and Doya, 2009).

At the same time, the ventral pallidum is also categorized as part of the basal forebrain circuitry (Zaborszky et al., 2012; Faget et al., 2018), integrating limbic and cognitive signals. Indeed, Avila and Lin found that putative GABAergic VP neurons with a bursting phenotype resemble those of other basal forebrain regions and share their salience-coding properties (Lin and Nicolelis, 2008), suggesting that salience information in the VP might be conveyed by afferents characteristic to the basal forebrain (Avila and Lin, 2014). In line with this, while reward-related signals in the VP were typically attributed to nucleus accumbens inputs (Kupchik et al., 2015; Root et al., 2015; Creed et al., 2016; Pardo-Garcia et al., 2019), a recent study (Ottenheimer et al., 2018) found earlier and stronger reward value signals in the VP when performing a direct comparison with the accumbens, raising the possibility that other afferents may play a major role in these rapid reward responses (Ottenheimer et al., 2018; Soares-Cunha and Heinsbroek, 2023). Another study found that somatostatin-expressing GABAergic VP neurons participate in controlling cortical gamma oscillations (Espinosa et al., 2019), which is a well-established function of the basal forebrain’s cortical projections, implicated in controlling attention and arousal (Kim et al., 2015; Yang et al., 2017; Király et al., 2023).

Interpreting the VP as a basal ganglia output has initially directed the focus to VP GABAergic neurons (van den Bos and Cools, 1991; Soares-Cunha and Heinsbroek, 2023); however, the VP contains considerable glutamatergic and cholinergic populations that have been addressed more recently (Faget et al., 2018; Stephenson-Jones et al., 2020; Farrell et al., 2021). Stephenson-Jones and colleagues found that both GABAergic and glutamatergic VP neurons can drive movement, but they are active in opposite valance contexts: GABAergic cells represent positive values and drive approach, while glutamatergic neurons represent negative values and drive avoidance (Stephenson-Jones et al., 2020). Similar results were found in the context of cocaine seeking (Heinsbroek et al., 2020), and it was shown that distinct inhibitory and excitatory VP projections mediate different aspects of depression-like symptoms (Knowland et al., 2017) and alcohol relapse (Prasad et al., 2020). Differential nucleus accumbens inputs to VP GABAergic vs. glutamatergic neurons were proposed to at least partially underlie the above differences (Neuhofer and Kalivas, 2023).

Comparably less is known about VP cholinergic neurons (Walaas and Fonnum, 1979; Zaborszky et al., 2012; Root et al., 2015). Cholinergic-specific VP lesions increased active coping mechanisms in fearful situations in mice (Akmese et al., 2023). In line with this, optogenetic stimulation of VP to basolateral amygdala cholinergic projections reduced pain thresholds and increased depression-like behaviors (Ji et al., 2023). Kim and colleagues found that this projection mostly codes aversive information, while a distinct set of cholinergic neurons represent appetitive cues in the context of odor discrimination (Kim et al., 2024).

However, a comprehensive account of the basal forebrain cholinergic population, including input-output mapping and their functional positioning along the basal ganglia - basal forebrain axis is missing, limiting the understanding of VP circuitry and functions. We fill this knowledge gap by revealing long range VPCN output connectivity including cortical projections characteristic of BFCNs and unlike striatal cholinergic interneurons and showing that VPCNs largely resemble BFCNs in their in vitro electrophysiological properties as well as in vivo responses to task-relevant salient stimuli.

## Materials and methods

### Animals

For targeted in vitro electrophysiological characterization of VPCNs and CINs, fluorophor expression in cholinergic neurons was driven by crossing ChAT-Cre and Ai32 (n = 4, 2/4 males, P40-60) or ChAT-Flp and Ai213 mice (n = 1 male, P40). ChAT-Cre mice were used for the characterisation of Reg- and Burst-BFCNs (n = 12, 7/12 males, p50-150). ChAT-Cre mice were used for anterograde (n = 6, 4/6 males, P50-100) and retrograde (n = 6, 3/6 males, P50-100) anatomical tracings. For the fiber photometry measurements, we used ChAT-Cre mice (n = 22, 14/22 males, P90-120). All experiments were conducted according to the regulations of the European Community’s Council Directive of 24 November 1986 (86/609/EEC); experimental procedures were reviewed and approved by the Animal Welfare Committee of the Institute of Experimental Medicine, Budapest and the Committee for Scientific Ethics of Animal Research of the National Food Chain Safety Office.

### Surgeries and viruses

The mice were anesthetized using a ketamine-xylazine solution (83 mg/kg ketamine and 17 mg/kg xylazine, prepared in 0.9% saline). After shaving and disinfecting the scalp with Betadine, the skin and subcutaneous tissues were numbed topically with Lidocaine spray. The mice were then positioned in a stereotaxic frame (Kopf Instruments), and their eyes were protected with Corneregel eye ointment (Bausch & Lomb). A sagittal incision was made in the skin using a surgical scalpel, exposing the skull, which was then cleaned. A craniotomy was drilled above the targeted area. For anterograde and retrograde tracings the craniotomy was opened above the ventral pallidum (VP, antero-posterior 0.5 mm; lateral 1 mm). Virus injections were performed for anterograde and retrograde tracing using a stereotaxic frame and a programmable nanoliter injector (Drummond Nanoject III). For anterograde tracing, AAV2/5.EF1a.Dio.hChR2(H134R)-eYFP.WPRE.hGH (Addgene; titer ≥ 1×10¹³ vg/mL) was injected into the VP at a dorso-ventral depth of 4.20 mm (20-30 nl). Retrograde tracing involved sequential injections of AAV2/9-Syn-FLEX-nGToG-WPRE3 (50 nl, Cat#BA-96, VCF of the Charité, Berlin) and, after a 4-week interval, pSADB19dG-mCherry (100 nl, Cat#BR-001, VCF of the Charité, Berlin) at the same dorso-ventral depth. For targeted in vitro electrophysiological characterization of Reg- and Burst-BFCNs, AAV2/5-EF1a-DIO-hChR2(H134R)-mCherry-WPRE-HGHpA was injected either into the caudal NB (antero-posterior −0.9 mm, lateral 2.2 mm, 3-4 dorso-ventral levels between 3.3 and 5 mm) or the HDB (antero-posterior 0.75 mm, lateral 0.6 mm, 2 dorso-ventral levels between 4.5 and 5.5 mm)(Laszlovszky et al., 2020). For fiber photometry experiments, mice were injected with AAVD7/2-CAG-hsyn-jGCaMP8m(rev)-dlox-WPRE-SV40r(A) (HDB and VP, 150 nL each side, HDB: antero-posterior 0.75 mm, lateral −0.60 mm; dorso-ventral −4.7 mm, VP: antero-posterior −0.61 mm, lateral 1.00 mm; dorso-ventral −4.5 mm). Anterograde virus injections were allowed a 4-week expression period, whereas retrograde tracings included a 9-day expression period following the rabies injection. During fiber photometry surgeries, injections were followed by the bilateral implantation of 400 μm core diameter optic fibers with ceramic ferrules (HDB, antero-posterior 0.75 mm, lateral −2.10 mm, dorso-ventral −4.5 mm; 20 degree lateral angle, VP, antero-posterior −0.61 mm, lateral 1.00 mm, dorso-ventral −4.3 mm; 0 degree lateral angle). The implant was secured to the skull with Super-Bond (Sun Medical Co.) and dental cement. Mice received analgesics (Buprenorphine, 0.1 mg/kg), local antibiotics (Gentamycin) and were allowed 10 days of recovery before starting behavioral training. All experiments were concluded by transcardial perfusion, and mouse brains were processed for further immunohistology experiments (see below).

### Behavioral training for fiber photometry

Mice were trained in a head-fixed auditory Pavlovian conditioning task using a probabilistic reinforcement schedule. The behavioral setup was custom-built to allow millisecond precision control of stimulus and reinforcement timing, as described in (Solari et al., 2018). Mice were subjected to a standard water restriction protocol prior to training and earned small water rewards (4 μL) during the conditioning phase. Two pure tones of different pitch (4 and 12 kHz, balanced across n = 10 and n = 12 mice; duration, 1s) predicted water reward or air-puff punishment with 90% probability (10% omissions). The two cue tones were presented in a randomized 50-50% ratio. All tones were set to 65 dB SPL. After the onset of the tone, mice could lick a waterspout, and individual licks were recorded by detecting when their tongues interrupted an infrared beam. Following a 400-600 ms post-stimulus delay, the scheduled outcome (water, air-puff, or omission) was delivered in pseudorandomized order based on the cue contingencies. Each new trial began after the animal refrained from licking for a minimum of 2.5 seconds. A foreperiod of 2.5–5.5 seconds, determined by a truncated exponential distribution, preceded each stimulus to prevent temporal expectations. Trials were restarted if the mouse licked during this foreperiod. Task control was handled by the Bpod behavioral system (Sanworks LLC, US). Air-puffs, 200 ms in duration, were delivered at 15 psi pressure, which stimulus was reported as aversive for head-fixed mice (Najafi et al., 2014; Hangya et al., 2015).

### Fiber photometry imaging

Dual-channel fiber photometry was used to monitor bilateral calcium activity, with fluorescence signals visualized throughout training sessions using the Doric Studio Software (Doric Neuroscience). Two LED sources (465 nm and 405 nm) were used in combination with fluorescent Mini Cubes (iFMC4, Doric Neuroscience). Amplitude modulation of the LEDs was achieved via a two-channel driver (LEDD_2, Doric Neuroscience), with 465 nm light modulated at 208 Hz and 405 nm light modulated at 572 Hz. The light was delivered to 400 µm patch cord fibers and connected to optical implants during the sessions. The same fibers were used to collect the fluorescence emitted from the tissue, which was detected by 500–550 nm photodetectors integrated into the Mini Cubes. Signals were sampled at 12 kHz, digitally decoded, and saved in *.csv format for later analysis.

### Perfusion

Mice were anesthetized with 2% isoflurane followed by an intraperitoneal injection of a mixture of ketamine-xylazine and promethazinium-chloride (83 mg/kg, 17 mg/kg and 8 mg/kg, respectively). After achieving deep anesthesia, mice were perfused transcardially (by placing the cannula into the ascending part of the aorta via an incision placed on the left ventricle wall) with saline for 2 minutes, followed by 4% paraformaldehyde (PFA) solution for 40 minutes, then saline for 10 minutes. After perfusion, mice were decapitated, and brains were carefully removed from the skull and postfixed in PFA overnight.

### Track verification for fiber photometry

A block containing the full extent of the HDB and VP was prepared, and 50 µm thick sections were cut using a Leica 2100S vibratome. All attempts were made to section parallel to the canonical coronal plane to aid track reconstruction efforts. All sections that contained the tracks were mounted on slides in Aquamount mounting medium. Epifluorescense images of the sections were taken with a Nikon C2 confocal microscope or Panoramic Midi Slidescanner. Atlas images were aligned to fluorescent images of the brain sections showing the fiber tracks and green fluorescent labeling in the target area. Only those recordings that were unequivocally localized to the HDB and VP were analyzed in this study.

### Anterograde and retrograde tracing

In case of anterograde tracing experiments, coronal sections of 50 µm thickness were cut by a vibratome (Leica VT1200S). Sections were extensively washed in 0.1M PB and TBS and blocked in 1% human serum albumin (HSA; Sigma-Aldrich) solution for 1 h. Then, sections were incubated in primary antibodies against eGFP (Thermo Fisher Scientific, Cat#A10262, 1:2000, raised in chicken; Table S2) for 48-60 hours. Sections were rinsed 3 times for 10 minutes in TBS; secondary fluorescent antibodies were applied overnight (anti-chicken Alexa-488, Jackson Immunoresearch, Cat#703-545-155, 1:1000; Table S3). Sections were rinsed in TBS and 0.1 M PB and mounted on slides in Aquamount mounting medium (BDH Chemicals Ltd). Sections containing the VP were incubated in primary antibody against ChAT (Synaptic Systems, Cat#297013, 1:500, raised in rabbit, Table S2), and anti-rabbit Alexa-594 secondary antibody (Thermo Fisher Scientific, Cat#A21207, 1:500, Table S3). Next, fluorescent images were taken with a Nikon A1R Confocal Laser Scanning Microscope. In the target areas of the VPCNs, fluorescent images were captured at 20x magnification using a standardized volume. Axon densities of VPCN projections were analyzed by classifying pixels with Ilastik (Berg et al., 2019). Axonal crossings were then quantified using a 5×5 grid overlay on the axon probability matrices to estimate axon densities.

In case of retrograde tracing experiments, coronal sections of 50 µm thickness were cut by a vibratome (Leica VT1200S). Sections were extensively washed in 0.1M PB and TBS and blocked in 1% human serum albumin (HSA; Sigma-Aldrich) solution for 1 h. Then, sections were incubated in primary antibodies against eGFP (Thermo Fisher Scientific, Cat#A10262, 1:2000, raised in chicken, Table S2) and mCherry (Biovision, Cat#5993-100, 1:1000, raised in rabbit, Table S2) for 48-60 hours. Sections were rinsed 3 times for 10 minutes in TBS; secondary fluorescent antibodies were applied overnight (anti-chicken Alexa-488, Jackson Immunoresearch, Cat#703-545-155, 1:1000, anti-rabbit Alexa-594, Termo Fisher Scientific, Cat#A21207, 1:500, Table S3). Sections were rinsed in TBS and 0.1 M PB and mounted on slides in Aquamount mounting medium (BDH Chemicals Ltd). Every second section was sampled to measure and estimate the number of transsynaptically labeled input cells using a Zeiss Axioplan2 epifluorescent microscope and a Pannoramic Digital Slide Scanner (3DHISTECH Kft., Hungary).

### Acute in vitro slice preparation

Mice were decapitated under deep isoflurane anesthesia, and the brains were rapidly removed and placed in ice-cold cutting solution, pre-carbogenated (95% O₂–5% CO₂) for at least 30 minutes before use. The cutting solution consisted of (in mM): 205 sucrose, 2.5 KCl, 26 NaHCO₃, 0.5 CaCl₂, 5 MgCl₂, 1.25 NaH₂PO₄, and 10 glucose. Coronal slices, 300 μm thick, were prepared using a Vibratome (Leica VT1200S). Following acute slice preparation, slices were transferred to an interface-type holding chamber for at least one hour of recovery. This chamber contained ACSF solution maintained at 35 °C, which gradually cooled to room temperature. The ACSF solution consisted of (in mM): 126 NaCl, 2.5 KCl, 26 NaHCO₃, 2 CaCl₂, 2 MgCl₂, 1.25 NaH₂PO₄, and 10 glucose, saturated with carbogen gas as described above.

### In vitro electrophysiology recordings

Recordings were performed under visual guidance using Nikon Eclipse FN1 microscope with infrared differential interference contrast (DIC) optics. The flow rate of the ACSF was 4–5 ml/min at 30–32°C (Supertech Instruments, Pecs, Hungary). Patch pipettes were pulled from borosilicate capillaries (with inner filament, thin-walled, outer diameter (OD) 1.5) with a PC-10 puller (Narishige, Tokyo, Japan). Pipette resistances were 3–6 MΩ when filled with intrapipette solution. The composition of the intracellular pipette solution was as follows (in mM): 54 d-gluconic acid potassium salt, 4 NaCl, 56 KCl, 20 Hepes, 0.1 EGTA, 10 phosphocreatine di(tris) salt, 2 ATP magnesium salt and 0.3 GTP sodium salt; with 0.2 % biocytin; adjusted to pH 7.3 using KOH and with osmolarity of ∼295 mOsm/l). Recordings were performed with a Multiclamp 700B amplifier (Molecular Devices, San Jose, US), digitized at 10 or 20 kHz with Digidata analog-digital interface (Molecular Devices), and recorded with pClamp11 Software suite (Molecular Devices). Cholinergic neurons expressing GFP or mOrange were visualized with the aid of LED light sources (Prizmatix Ltd., Holon, Israel) integrated into the optical light path of the microscope and detected with a CCD camera (Andor Zyla). We applied a somatic current injection protocol containing a 3-s-long, incremental ‘prepolarization’ step followed by a positive square pulse (1 s), to elicit spiking starting from different membrane potentials similarly as in (Laszlovszky et al., 2020). Furthermore, we applied a simple step protocol consisting of a series of hyperpolarizing and depolarizing steps, each lasting 1 second, to further determine the spiking characteristics of distinct cholinergic cell types.

### Immunohistochemical identification of in vitro recorded cholinergic cells

After acute slice electrophysiology experiments, brain sections were fixed overnight in 4% PFA. Sections were extensively washed in 0.1M PB and TBS and blocked in 1% human serum albumin (HSA; Sigma-Aldrich) solution for 1 h. Then, sections were incubated in primary antibodiy against ChAT (Synaptic Systems, Cat#297013, 1:500, Table S2) for 48-60 hours. This step was followed by thorough rinse with TBS (3 × 10 minutes) and overnight incubation with a mixture of anti-rabbit Alexa-594 secondary antibody (Thermo Fisher Scientific, Cat#A21207, 1:500, Table S3) and streptavidin-A488 (Invitrogen, Cat#S11223, 1:1000). We used 0.1% Triton-X detergent through every incubation step due to the thickness of the brain section. Finally, sections were washed in TBS and PB, mounted on microscopy slides, covered with Vectashield (Vector Laboratories Inc, US) and imaged with a Nikon A1R confocal laser scanning microscope.

### Analysis of in vitro experiments

All in vitro data were processed and analyzed offline using Python 3. Spike delay was defined as the interval between the start of the 1-second positive current injection step and the peak time of the first action potential (AP) and was calculated using the ’prepolarization’ protocol. Burst frequency was determined from the subsequent three inter-spike intervals (ISIs). Autocorrelograms (ACGs) for each cell were computed using spikes evoked by simple step protocols and were smoothed with a 5-ms moving average. Electrophysiological features were extracted using the Electrophys Feature Extraction Library (eFEL, (Ranjan et al., 2024)). These features were derived from the first current injection step that elicited at least four action potentials in simple step protocols. The resulting dataset was used for the low dimensional projection with Uniform Manifold Approximation (UMAP, (McInnes et al., 2018)).

### Analysis of in vivo electrophysiology data

Data were processed in Matlab R2018a. Auto-correlograms (ACG) were computed at 0.5 ms resolution and smoothed using a 2.5 ms (5-point) moving average for visualization. Individual ACGs were normalized to their mean values, sorted by Burst Index or refractory period, and averaged per group. The Burst Index was calculated as the difference between the maximum ACG at 0-25 ms and the mean ACG at 180-200 ms, normalized by the larger of these two values, yielding an index between −1 and 1 (modified from (Laszlovszky et al., 2020) based on the slower bursts of VPCNs). The Theta Index was calculated based on the difference between the mean ACG values within a ±25 ms window around the 5-10 Hz theta peak (100-200 ms lags). Refractory periods were estimated by identifying low-probability spiking intervals from the ACGs, using a 10 ms moving average to find the half-height point of the ACG’s central trough. This provided a measure of relative refractory periods rather than absolute spike repolarization (Royer et al., 2012; Laszlovszky et al., 2020).

### Analysis of fiber photometry recordings

Matlab R2018a (Mathworks, Natick) was used to process fiber photometry data, following the procedures described in refs. (Lerner et al., 2015; Hegedüs et al., 2023). Animals with sufficient viral expression in the target region, as well as successful surgical targeting that resulted in measurable fluorescent signals were included in the analyses, resulting in n = 21 mice for VP-specific analyses, n = 16 mice for HDB-specific analyses, and n = 15 mice for VP-HDB comparisons. The fluorescence signals were digitally filtered below 20 Hz using a low-pass Butterworth filter to remove high-frequency noise. The delta fluorescence (dF/F) signal was computed by fitting a least-squares regression to the 405 nm isosbestic control signal and aligning its baseline with that of the 465 nm calcium-dependent signal (f465). The normalized 405 nm signal (f405,fitted) was then subtracted from the 465 nm signal as follows: dF/F = (f465 - f405,fitted) / f405,fitted * 100, to account for motion artifacts and autofluorescence. Slow baseline decay was corrected with a 0.2 Hz high-pass filter. The dF/F signals were Z-scored relative to the mean and standard deviation of a baseline period (2 seconds before cue onset) and averaged across trials. The analysis included only the last 5 sessions where Cue 1 and Cue 2 occurred with equal (0.5) probabilities. Response maxima, along with latency, duration, and area under the curve (AUC), were computed. Cross-correlations (CCG) were computed between two photometry signals (VP and HDB) at the maximal time resolution allowed by the sampling rate (12048 Hz) using the built-in xcorr function in MATLAB. To reject common mode noise, the central ±20 ms window around 0-ms lag was excluded from the analysis.

### Analysis of pupil dynamics

To monitor pupil dynamics during behavioral training, we used a Flea3 FL3-U3-32S2M camera focused on the mouse’s eye. Video capture was synchronized with the fiber-photometry recording through TTL signals, with a TTL pulse sent at the beginning of each frame and recorded at 59 FPS. The videos were analyzed offline using DeepLabCut (Mathis et al., 2018), which was trained to track pupil edges at three diagonal points and eyelid positions. Pupil diameter was calculated as the mean distance between the three diagonal points and interpolated to match the sampling rate of the fiber-photometry data. Calcium transient peaks recorded in either the ventral pallidum (VP) or the horizontal diagonal band of Broca (HDB) were used to calculate VP/HDB activity-evoked changes in pupil size (Figure 7D). Transfer entropy values were computed on the z-scored, downsampled, and discretized VP/HDB and pupil time series using the PyInform Python library for information-theoretic measures of time-series data. During discretization, the continuous data were divided into 200 equally spaced bins, and each data point was assigned to its corresponding bin.

### Statistical Analysis

We estimated the sample size before conducting the study based on previous publications (Laszlovszky et al., 2020; Hegedüs et al., 2023) and corresponding statistical power estimations (https://github.com/hangyabalazs/statistical-power). The study did not involve separate experimental groups, so randomization and blinding were not relevant to the study. Automated data analysis was conducted independently of neuron identity. For neurons with more than 50,000 spikes, ACG calculation was capped at 50,000 spikes to avoid memory limitations. Comparisons between conditions were performed using non-parametric tests to avoid assumptions on normality, which could not be confirmed statistically. The Wilcoxon signed-rank test was used for paired samples, while the Mann-Whitney U-test was used for unpaired comparisons. Peri-event time histograms (PETHs) were presented as mean ± SE, while box plots showed median, interquartile range, and non-outlier range, with all data points displayed.

## Results

### Input-output connectivity of VPCNs reveal broad connections with the affective circuit

Mapping of afferent and efferent connectivity of BFCNs have been carried out for the broadly defined basal forebrain (Do et al., 2016; Hu et al., 2016); however, these experiments did not include specific VP injections. Additionally, cholinergic output connectivity was determined for the SI, HDB and MS regions (Saper, 1984; Zaborszky et al., 2012; Agostinelli et al., 2019) but not for the VP.

To fill this gap, we first performed anterograde tracing of VPCNs by injecting AAV2/5-EF1a-DIO-EYFP in the VP region of ChAT-Cre mice. We found that VPCNs projected robustly to the basolateral amygdala, nucleus accumbens, prefrontal cortex, and, to lesser extent, to the lateral habenula and the parasubthalamic nucleus. This projection pattern was concordant with general BFCN projections to ventral striatum, prefrontal cortex and amygdala (Do et al., 2016), but differed from specific subcortical projections of the horizontal nucleus of the diagonal band of Broca (HDB), the medial septum (MS) and the substantia innominata (SI) cholinergic neurons in the absence of hippocampal but presence of nucleus accumbens targets (Agostinelli et al., 2019).

Next, we performed input mapping of VPCNs by mono-transsynaptic rabies tracing. We found that VPCNs received the majority of their monosynaptic inputs from the nucleus accumbens, the lateral hypothalamus, and the central amygdala, with smaller contributions from the preoptic area, and the bed nucleus of stria terminalis. This afferent connectivity aligns with previously reported inputs to BFCNs (Do et al., 2016; Hu et al., 2016).

### Ventral pallidal cholinergic neurons resemble the bursting type of basal forebrain cholinergic neurons

BFCNs form two distinct cell types, a synchronous population of neurons that fire bursts that correlate with cortical activity (Burst-BFCNs), and a regular rhythmic firing group of cells that synchronizes with cortical activity in a behavior-predictive manner (Reg-BFCN) (Laszlovszky et al., 2020; Lozovaya et al., 2024). Striatal cholinergic interneurons (CINs) are relatively homogeneous and their firing pattern resembles that of Reg-BFCNs (Inokawa et al., 2010; Zhang et al., 2018; Cox and Witten, 2019). We characterized intrinsic electrophysiological properties of VPCNs, BFCNs and CINs with identical protocols to determine how VPCN activity is related to the above better-known cholinergic populations.

We prepared acute slices from mice expressing fluorescent proteins selectively in cholinergic neurons (see Methods) and performed whole-cell patch clamp recordings from n = 20 VPCNs (Fig.3A). These recordings were contrasted to novel acute slice recordings of CINs (n = 8) as well as previously obtained traces (Laszlovszky et al., 2020) of Burst-BFCNs and Reg-BFCNs (n = 29 and 31, respectively; Fig.3B-C). Autocorrelations of VPCN activity during somatic current injection protocols revealed a homogeneous bursting phenotype, resembling burst-BFCNs of the HDB and SI (Fig.3D). However, they were differentiated from the latter group by somewhat longer refractory periods and lower maximal burst frequency (Fig.3E-H). In sum, VPCNs form a distinct group based in their intrinsic electrophysiological properties, closely resembling burst-BFCNs of the HDB and SI.

**Figure 1.**
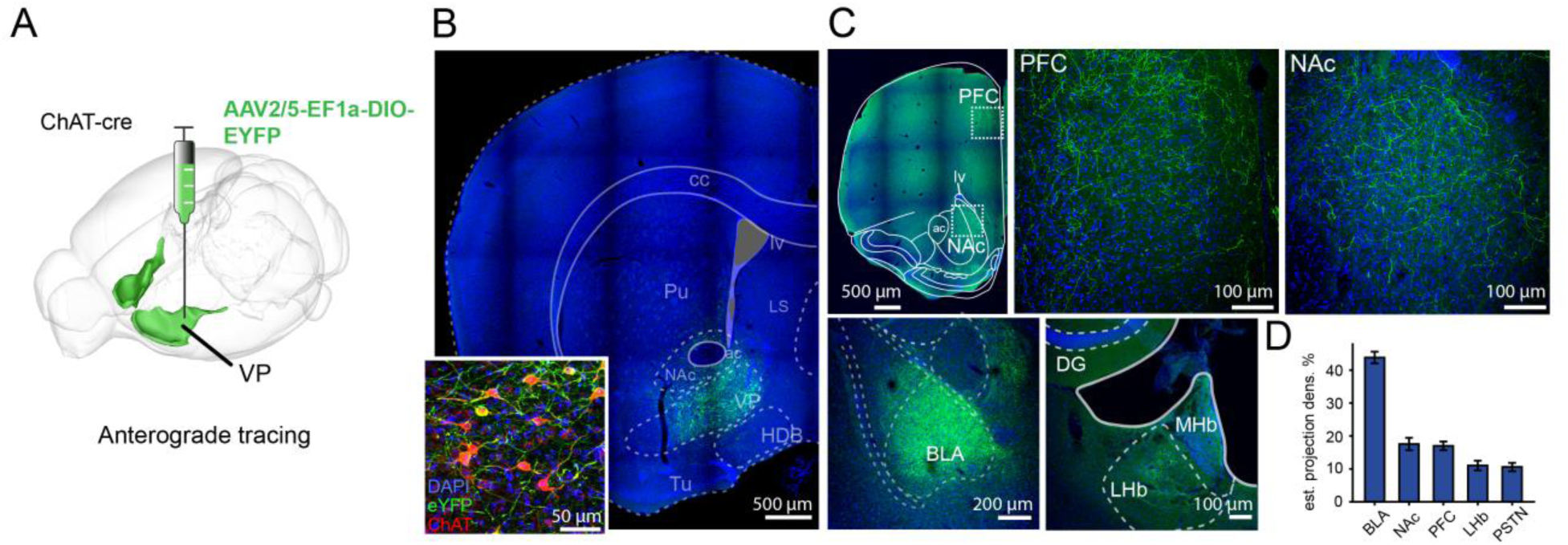
VPCNs innervate the affective brain circuitry. **(A)** Schematic illustration of an anterograde tracer virus injection into the ventral pallidum of a ChAT-cre mouse. **(B**) Fluorescent image of the injection site, showing eYFP (green) and DAPI (blue) labeling. **(C)** Fluorescent images showing the main target areas innervated by VPCNs, including the prefrontal cortex (PFC), the nucleus accumbens (NAc), the basolateral amygdala (BLA), and the lateral habenula (LHb); green, eYFP; blue, DAPI. **(D)** Estimated projection density in the primary output regions of VPCNs, expressed as a percentage of labeled axons (n = 6 mice). Bars and error bars represent mean ± SEM (BLA, 43.81 ± 1.77%; NAc,17.56 ± 1.88%; PFC,17.00 ± 1.33%; LH,11.04 ± 1.47%; PSTN,10.59 ± 1.26%).

**Figure 2.**
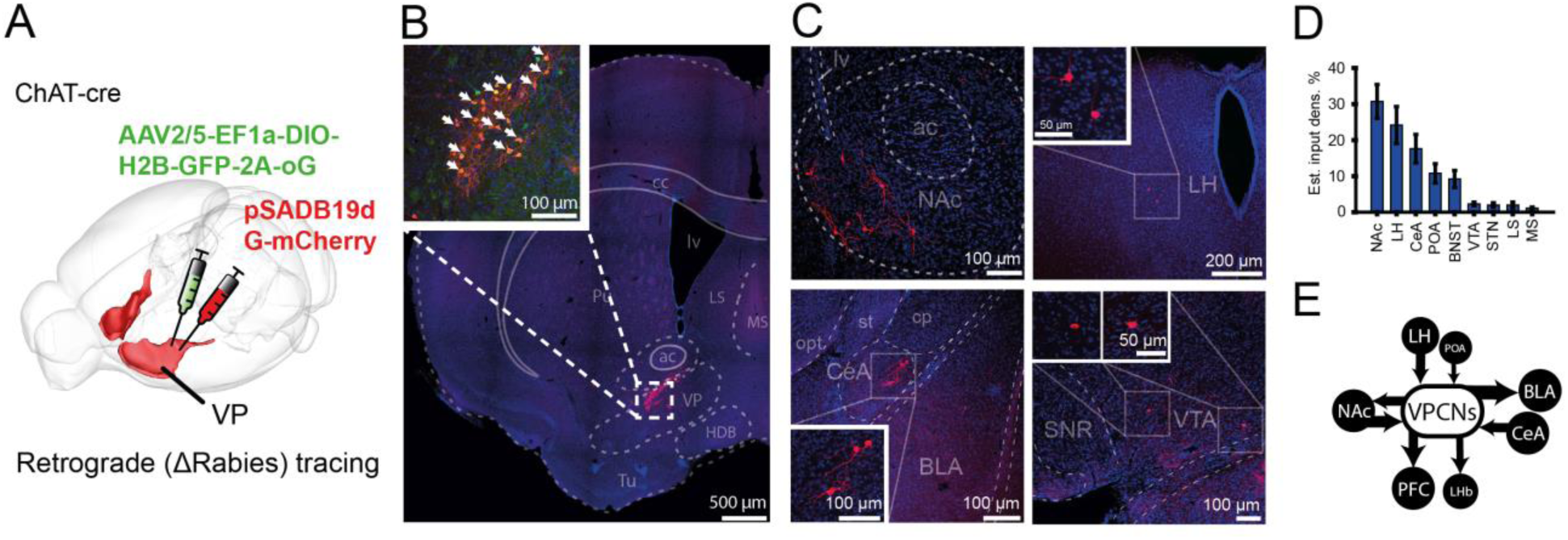
VPCNs receive inputs from the affective brain circuitry. **(A)** Schematic illustration of the injection site in the ventral pallidum of a ChAT-cre mouse, showing the delivery of the helper virus (green) and pseudotyped rabies virus (red). **(B)** Fluorescent image of the injection site. Inset, cells co-expressing the helper and rabies viruses (white arrowheads). **(C)** Fluorescent images showing input cells in the nucleus accumbens, the lateral hypothalamus, the central amygdala, and the ventral tegmental area. **(D)** Estimated input density as a percentage of total input cells (n = 703 cells from 6 mice) across various brain regions. Bars and error bars represent mean ± SEM (NAc, 30.73 ± 4.69%; LH, 24.18 ± 5.14%; CeA, 17.64 ± 3.94%; POA, 10.81 ± 2.67%; BNST, 9.25 ± 2.36%; VTA, 2.28 ± 0.48%; STN, 1.99 ± 0.62%; LS, 1.99 ± 0.79%; MS,1.14 ± 0.28%). **(E)** Schematic summary of the major input and output regions of the VPCNs.

**Figure 3.**
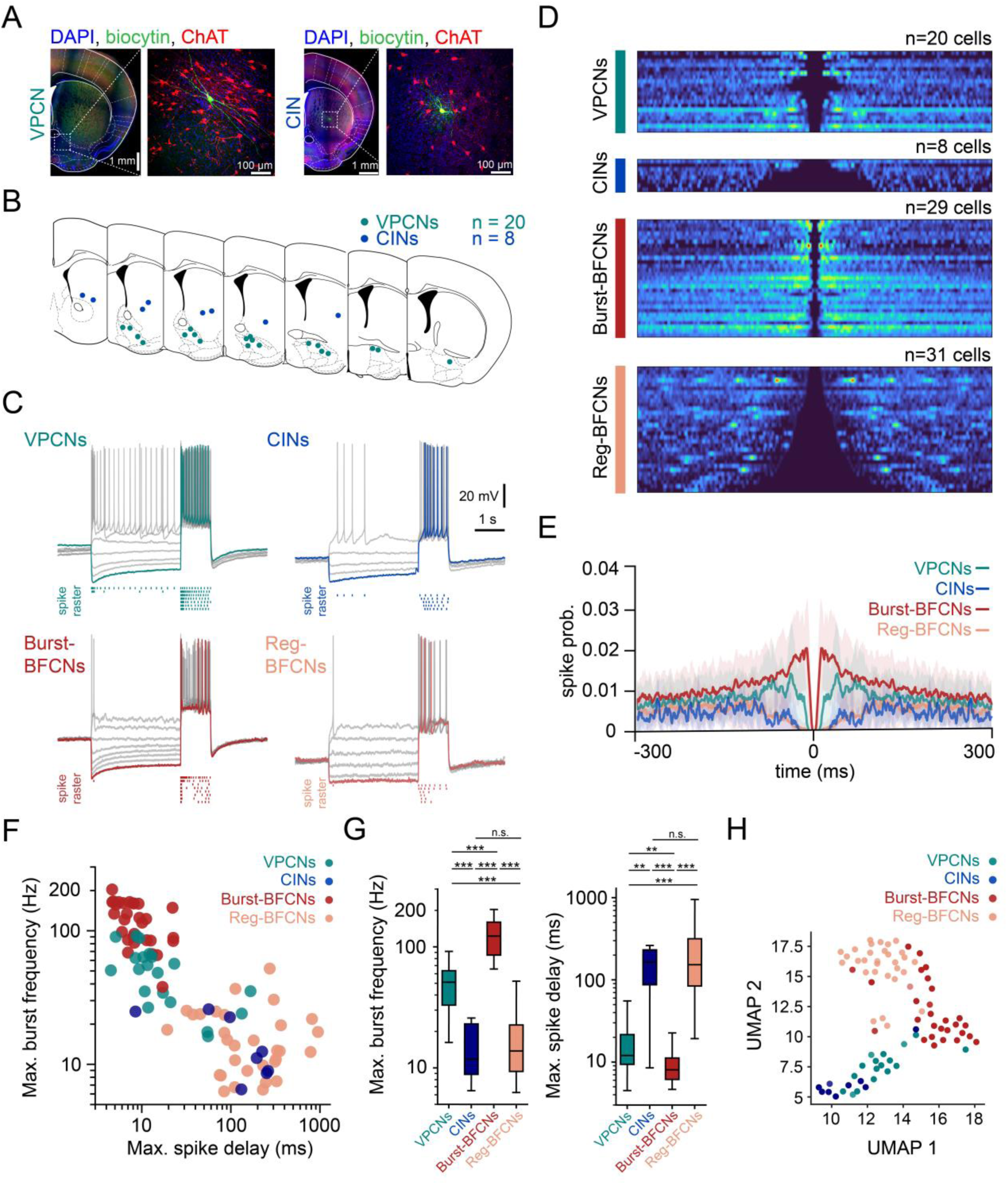
VPCNs resemble burst-firing cholinergic cells of the basal forebrain. **(A)** Representative confocal images of a recorded and biocytin-filled (green) VPCN (left; scale bars, 1 and 0.1, mm respectively) and a CIN (right; scale bars, 1 and 0.1 mm, respectively) from a reporter mouse expressing red fluorescent protein in all cholinergic neurons. **(B)** Locations of the recorded VPCNs (n = 20) and CINs (n = 8). **(C)** Representative firing patterns of ventral pallidal, striatal, basal forebrain burst-, and regular-firing cholinergic cells. VPCNs display short spike delays and high-frequency spike clusters in response to positive current injections, resembling burst-firing basal forebrain cholinergic neurons (Burst-BFCNs). In contrast, CINs exhibit firing patterns similar to regular-firing BFCNs (Reg-BFCNs). **(D)** Spike autocorrelograms during somatic current injection protocols for all recorded cholinergic neurons, grouped by cell type. **(E)** Average autocorrelograms for VPCNs (teal, n = 20), CINs (blue, n = 8), Burst-BFCNs (red, n = 29), and Reg-BFCNs (pink, n = 31). Solid lines represent the mean, and shaded regions indicate SEM. **(F)** Maximal burst frequency plotted against maximal spike delay for all recorded cells on a log-log scale, color-coded by cell type. **(G)** Population statistics comparing the maximum burst frequency and spike delay across all cholinergic neuron types. **, p < 0.01; ***, p < 0.001; Mann-Whitney U-test. Maximal spike delay, VPCNs vs. CINs, p = 0.0044; VPCNs vs. Burst-BFCNs, p = 0.00298; VPCNs vs. Reg-BFCNs, p = 2.21 × 10^−7^; CINs vs. Burst-BFCNs, p = 0.00012; CINs vs. Reg-BFCNs, p = 0.50502; Burst-BFCNs vs. Reg-BFCNs, p = 2.08 × 10^−11^. Maximal burst frequency, VPCNs vs. CINs, p = 4.31 × 10^−5^; VPCNs vs. Burst-BFCNs, p = 1.16 × 10^−7^; VPCNs vs. Reg-BFCNs, p = 3.34 × 10^−7^; CINs vs. Burst-BFCNs, p = 2.021 × 10^−5^; CINs vs. Reg-BFCNs, p = 0.88; Burst-BFCNs vs. Reg-BFCNs, p = 1.54 × 10^−11^. (**H**) Uniform Manifold Approximation and Projection (UMAP) of a high-dimensional electrophysiological feature set extracted from all cholinergic cells (see Methods), color-coded by cell type.

### VPCNs respond to rewards, punishments and reward-predicting cues

To determine the behavioral correlates of VPCNs, we trained head-fixed mice on Pavlovian conditioning, where two pure tones of different pitch (Cue1 and Cue2) predicted water reward or air puff punishment, respectively (Fig.4A). Mice learned these task contingencies, indicated by preferential anticipatory licking after the reward-predicting tone (Fig.4B-C).

**Figure 4.**
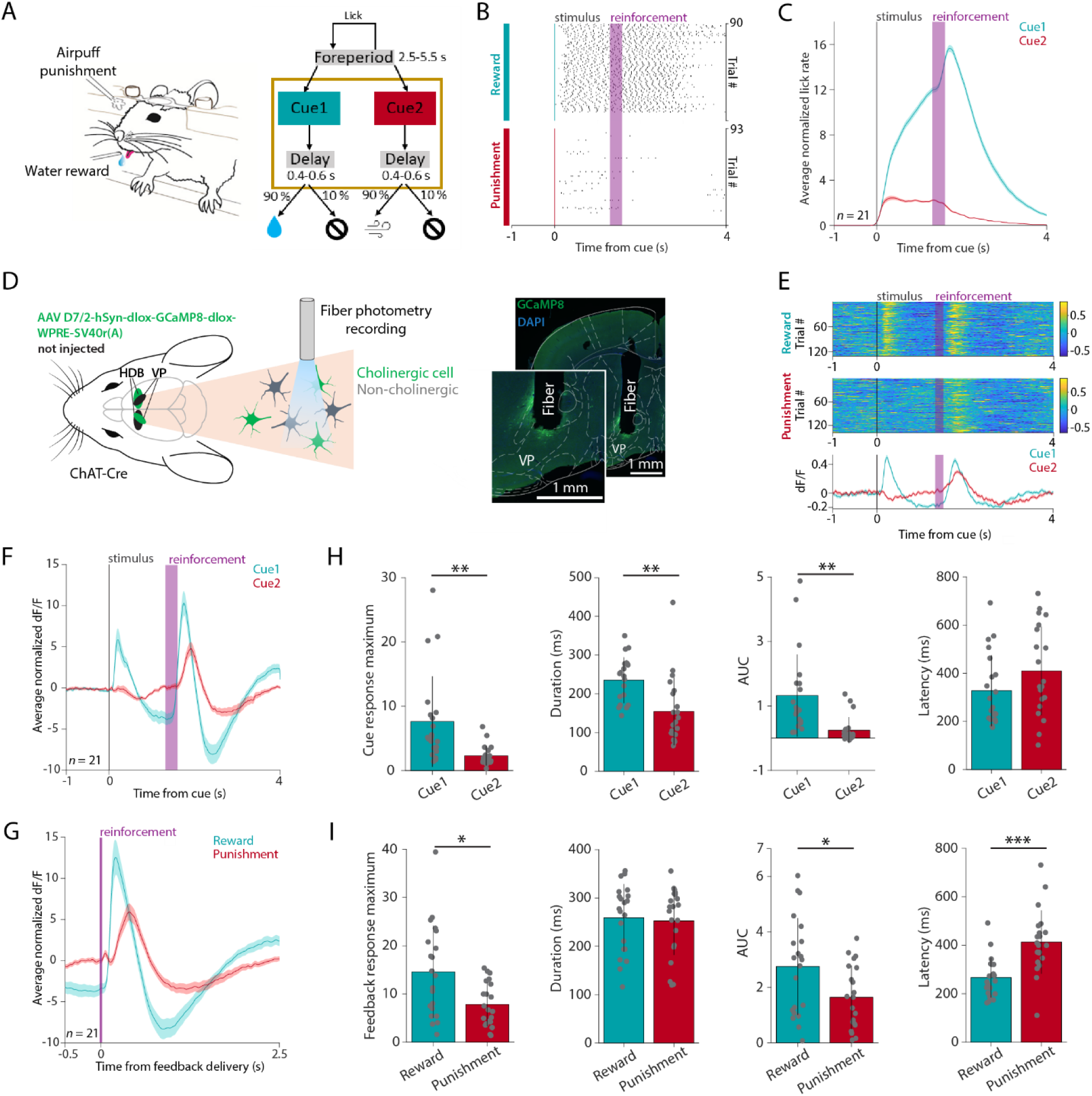
Cholinergic cells in the ventral pallidum respond differently to the reward- and punishment-predicting cues. **(A)** Schematic of the head-fixed probabilistic Pavlovian conditioning task. Created using Mathis, M. (2020), Classical Conditioning Mouse, Zenodo, https://doi.org/10.5281/zenodo.3925907, under Creative Commons 4.0 license (https://creativecommons.org/licenses/by/4.0/). The original image was not modified. **(B)** Raster plot of induvial licks aligned to the onset of Cue 1 and Cue 2, respectively, in an example recording session. The mouse showed preferential anticipatory licking to the reward-predicting Cue 1. **(C)** Average z-scored anticipatory lick rate of all animals (n = 21), aligned to the reward-predicting Cue 1 (green) and the punishment-predicting Cue 2 (red). Error shades indicate SEM. **(D)** Left, schematic representation of the fiber photometry measurements. We injected AAV D7/2-hSyn-dlox-GCaMP8-dlox-WPRE-SV40r(A) into the ventral pallidum (VP) and the horizontal limb of the diagonal band of Broca (HDB) in the two hemispheres of ChAT-Cre mice, and measured cholinergic calcium signals using fiber photometry. Created using Petrucco, L. (2020), Mouse head schema, Zenodo, https://doi.org/10.5281/zenodo.3925902 and Scidraw, S. (2020), Neuron silhouette, Zenodo, https://doi.org/10.5281/zenodo.3925927, under Creative Commons 4.0 license (https://creativecommons.org/licenses/by/4.0/). The original image was not modified. Right, representative fluorescent histological image of the measurement site (green, GCaMP8; blue, DAPI nuclear staining). Scale bars, 1 mm. **(E)** Example fiber photometry recording of VPCNs from a single session. Top, normalized dF/F traces of all rewarded and punished trials aligned to cue onset, color coded (blue, low values; yellow, high values). Bottom, average dF/F traces from the same session. Error shades indicate SEM. **(F)** Average z-scored dF/F of VPCNs aligned to the reward-predicting Cue 1 (green) and the punishment-predicting Cue 2 (red), averaged across all animals (n = 21). Error shades indicate SEM. **(G)** The same as in panel F but aligned to reward (green) and punishment delivery (red). Error shades indicate SEM. **(H)** From left to right, comparison of response magnitude, duration, integral and latency between VPCN responses to the reward-predicting Cue 1and the punishment-predicting Cue 2. Each dot represents the session-average of a single animal. AUC, area under the curve. Bar graphs show mean ± standard deviation. *, p < 0.05; **, p < 0.01; ***, p < 0.001; Wilcoxon signed-rank test. (**I**) The same as in panel H but comparing VPCN responses to reward and punishment. Each dot represents the session-average of a single animal. Bar graphs show mean ± standard deviation. *, p < 0.05; **, p < 0.01; ***, p < 0.001; Wilcoxon signed-rank test.

ChAT-Cre mice (n = 21) were injected with AAV D7/2-hSyn-dlox-GCaMP8-dlox-WPRE-SV40r(A) to express the fluorescent calcium indicator GCaMP8 in VPCNs and implanted with optic fibers in the VP and HDB regions on the two sides (Fig.4D). We performed fiber photometry recordings of bulk calcium levels of VPCNs while mice performed the Pavlovian task. We found that both VPCNs and HDB BFCNs consistently responded to cues, rewards and punishments (Fig.4E-G; Fig.S1). While the reward-predicting Cue1 evoked large increases in calcium, the punishment-predicting Cue2 induced smaller and more variable responses including an earlier increase and a later decrease (Fig.4G). Overall, responses to Cue1 were significantly larger in amplitude and integral, and longer in duration (Fig.4H).

Both rewards and punishments evoked consistent, large increases in VPCN calcium signals. A quantitative comparison revealed that reward responses were larger and faster than punishment responses (Fig.4I).

Next, we directly compared VPCN calcium responses to parallel recordings from the HDB of the basal forebrain (Fig.5). We found that the response patterns were qualitatively similar, including cue, reward and punishment responses (Hegedüs et al., 2023). The similarities between VPCN and BFCN response were not restricted to these signal correlations but were accompanied by consistent positive moment-by-moment noise correlations revealed by cross-correlation analysis, showing a zero-lag positive correlation flanked by negative correlations around a characteristic delay of approximately 0.8 s (−0.721 s and 0.861 s; Fig.5A-B). However, a quantitative comparison uncovered notable differences as well: while responses to the reward-predicting Cue1 were almost identical (Fig.5C,E), HDB BFCNs exhibited larger responses to the punishment-predicting Cue2 (Fig.5D,F). While reward-responses were much larger, longer and faster in VP (Fig.5G,I), the punishment responses were comparable in amplitude but faster in the HDB (Fig.5H,J). These results reveal BFCN-like response patterns in VPCNs, but with a quantitative preference to appetitive compared to aversive stimuli.

**Figure 5.**
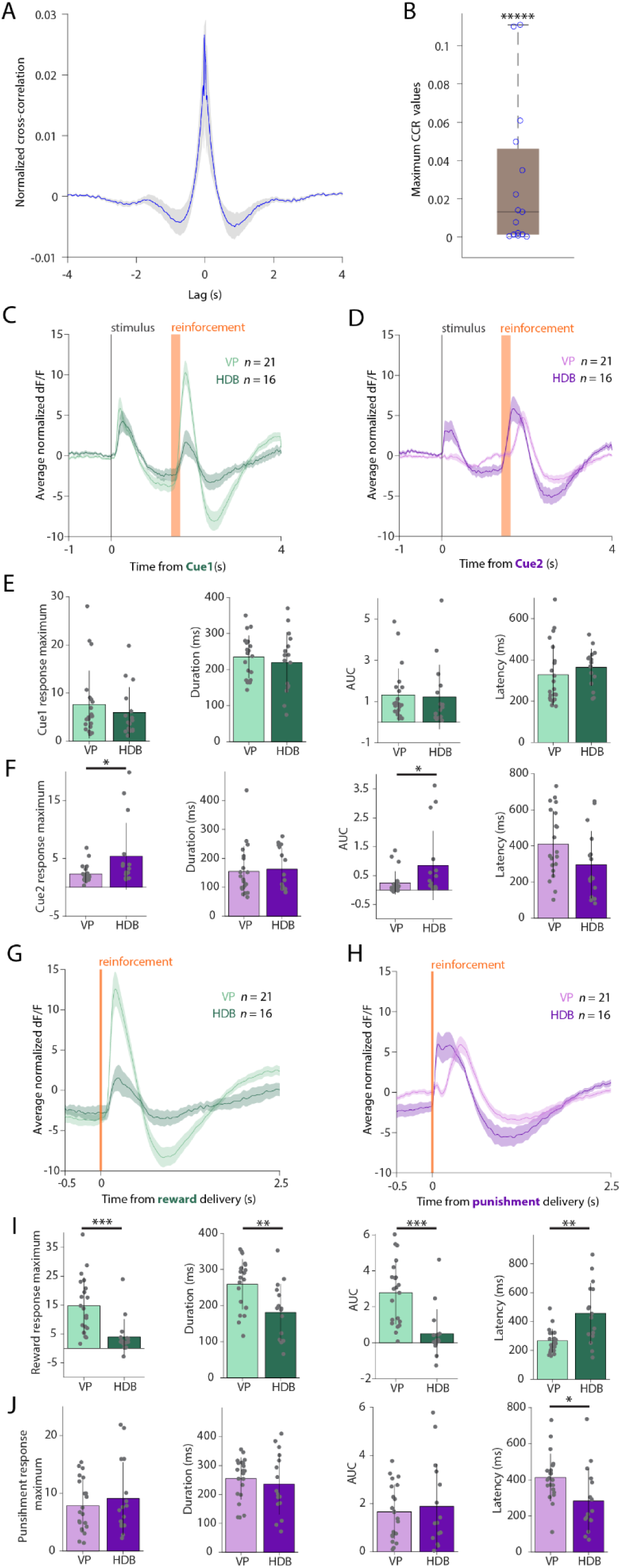
Differences of cholinergic reward and punishment responses between VP and HDB. **(A)** Cross-correlation of HDBCN and VPCN bulk calcium recordings, averaged across all animals (n = 15 mice with both signals accepted, see Methods). **(B)** Maximal cross-correlation (CCR) values averaged per mice (n = x). *****, p < 0.00001 for CCR > 0, Wilcoxon signed-rank test. **(C)** Average z-scored dF/F of VPCNs (light green) and HDBCNs (dark green) aligned to the reward-predicting cues, averaged across all animals (VP, n = 21; HDB, n = 16). Error shades indicate SEM. **(D)** Average z-scored dF/F of VPCNs (light purple) and HDBCNs (deep purple) aligned to the punishment-predicting cues, averaged across all animals (VP, n = 21; HDB, n = 16). Error shades indicate SEM. **(E)** From left to right, comparison of Cue1 response magnitude, duration, integral and latency between VPCNs (n = 21) and HDBCNs (n = 16). Each dot represents the session-average of a single animal. Bar graphs show mean ± standard deviation. Wilcoxon signed-rank test. **(F)** From left to right, comparison of Cue2 response magnitude, duration, integral and latency between VPCNs (n = 21) and HDBCNs (n = 16). Each dot represents the session-average of a single animal. Bar graphs show mean ± standard deviation. *, p < 0.05; Wilcoxon signed-rank test. **(G)** Average z-scored dF/F of VPCNs (light green) and HDBCNs (dark green) aligned to rewards, averaged across all animals (VP, n = 21; HDB, n = 16). Error shades indicate SEM. **(H)** Average z-scored dF/F of VPCNs (light purple) and HDBCNs (deep purple) aligned to punishments, averaged across all animals (VP, n = 21; HDB, n = 16). Error shades indicate SEM. **(I)** From left to right, comparison of reward response magnitude, duration, integral and latency between VPCNs (n = 21) and HDBCNs (n = 16). Each dot represents the session-average of a single animal. Bar graphs show mean ± standard deviation. **, p < 0.01; ***, p < 0.001; Wilcoxon signed-rank test. **(J)** From left to right, comparison of punishment response magnitude, duration, integral and latency between VPCNs (n = 21) and HDBCNs (n = 16). Each dot represents the session-average of a single animal. Bar graphs show mean ± standard deviation. *, p < 0.05; Wilcoxon signed-rank test.

### Most putative VPCNs show spike responses to salient stimuli like those of BFCNs

To assess the spiking heterogeneity of VPCNs corresponding to these bulk calcium responses, we analyzed the activity of putative VPCNs (pVPCNs) recorded in a similar Pavlovian conditioning task. In this task, different auditory cues predicted large water reward, small reward or no reward, large air puff punishment, small punishment or no punishment in blocks of positive and negative valence trials (Stephenson-Jones et al., 2020) (Fig.6A). VP neurons (n = 331 from 6 mice) fell in four distinct response categories by hierarchical clustering, and optogenetic tagging of glutamatergic and GABAergic neurons unambiguously identified two clusters as GABAergic and one as glutamatergic. The remaining ‘Type I’ neurons (n = 22 / 331) did not contain any glutamatergic or GABAergic neurons and were therefore identified as pVPCNs.

**Figure 6.**
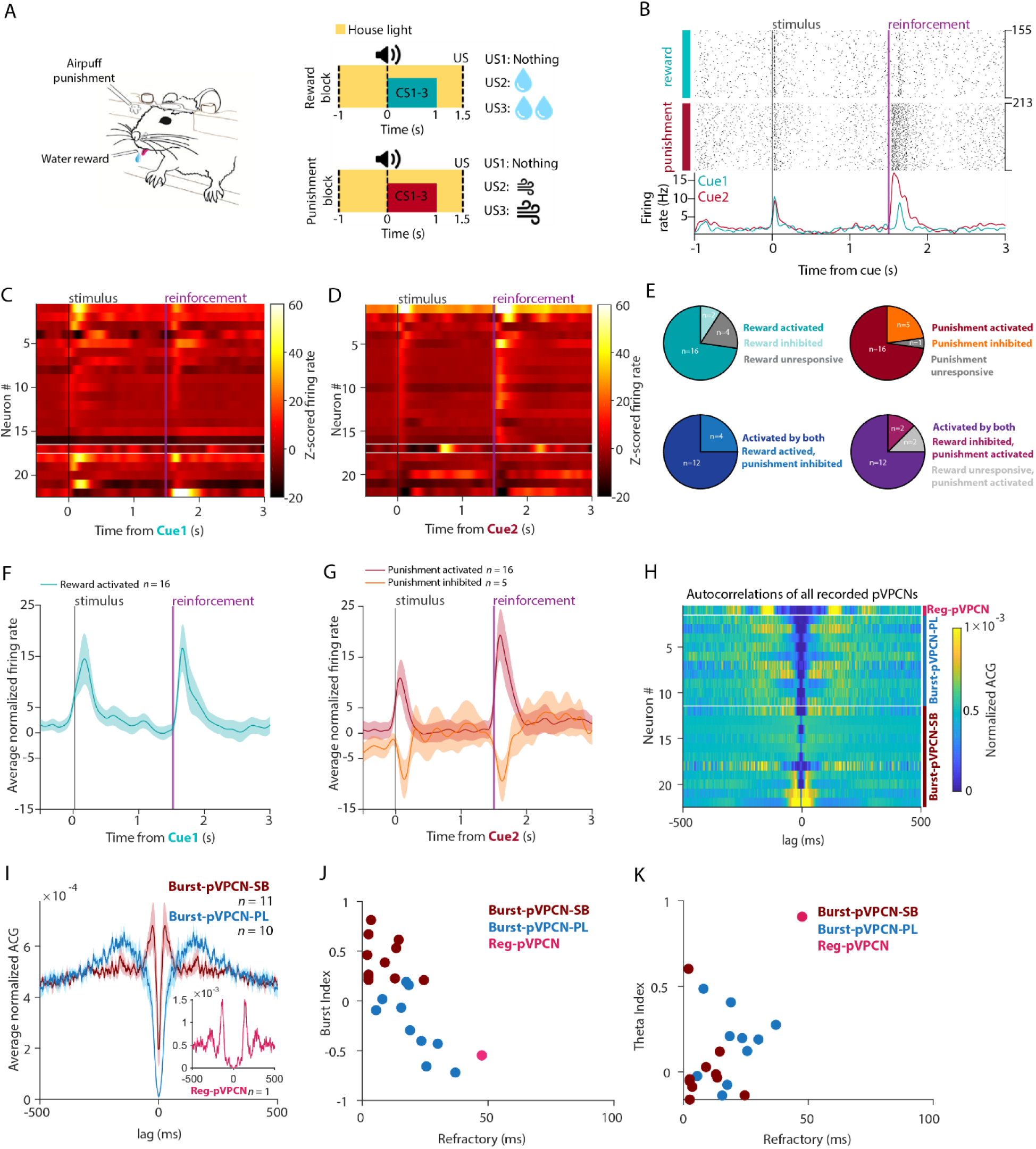
Most putative VPCNs show spike responses to salient stimuli like those of BFCNs. **(A)** Schematic of the Pavlovian conditioning task. **(B)** Top, raster plot of spike times aligned to cue onset of an example pVPCN during the Pavlovian task in rewarded and punished trials. Bottom, corresponding PETHs (green, rewarded trials; red, punished trials). **(C)** Z-scored PETHs of all recorded pVPCNs (n = 22) during rewarded trials shown on a heatmap, sorted by the punishment response magnitudes (for consistent ordering across panels C and D). Cells activated significantly after punishment are shown above the upper white line, while those that were significantly inhibited by punishment are shown below the lower white line. **(D)** Z-scored PETHs of all recorded pVPCNs (n = 22) during punishment trials shown as a heatmap. Cells activated significantly after punishment are shown above the upper white line, while those that were significantly inhibited are shown below the lower white line. **(E)** Pie charts showing the proportions of pVPCN response types. Top left, reward responses of all recorded pVPCNs (n = 22). Top right, punishment responses of all pVPCNs. Bottom left, punishment responses of reward-activated pVPCNs (n=16). Bottom right, reward responses of punishment-activated pVPCNs (n = 16). **(F)** The average normalized firing rate of reward-activated pVPCNs (n = 16) aligned to cue onset during rewarded trials (mean ± SEM). **(G)** The average normalized firing rate of punishment-activated pVPCNs (n = 16, dark red) and punishment-inhibited pVPCNs (n = 5, light red) aligned to cue onset during punishment trials (mean ± SEM). **(H)** Autocorrelations (ACG, normalized to a sum of one) of all recorded pVPCNs (n = 22). Reg-pVPCN, above the upper white line; Burst-pVPCN-PLs (n = 10), between the two horizontal white lines; Burst-pVPCN-SBs (n = 11), below the lower white line; sorted by Burst index within each group. **(I)** Normalized autocorrelation of all recorded pVPCNs averaged by firing pattern type. Dark red, Burst-pVPCN-SB (n = 11); blue, Burst-pVPCN-PL (n = 10), light red in inset, Reg-pVPCN (n = 1). **(J)** Burst Index vs. Refractory period of pVPCNs, color coded by firing pattern type. **(K)** Theta Index vs. Refractory period of pVPCNs, color coded by firing pattern type.

**Figure 7.**
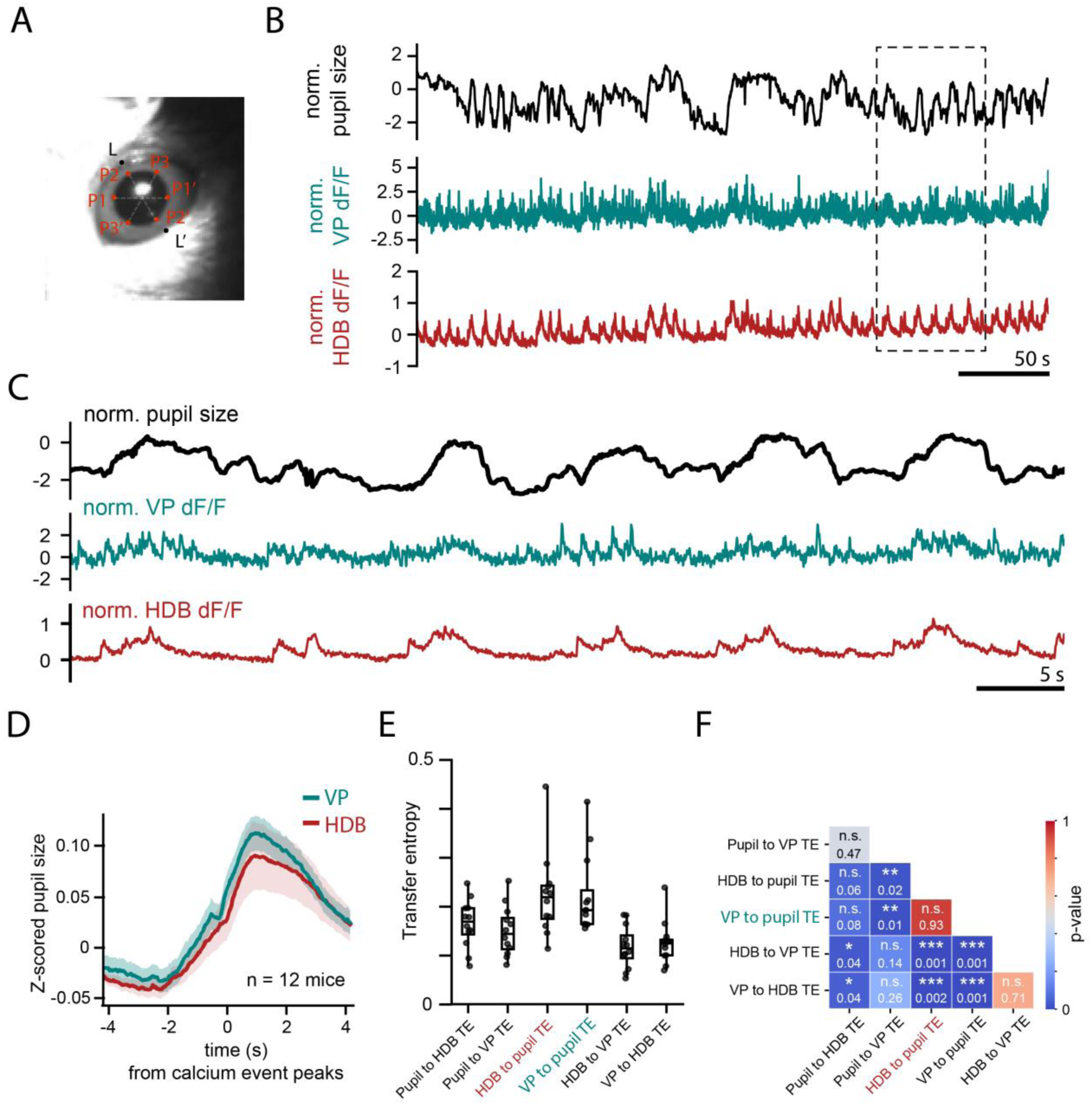
Pupil size correlates with both VPCN and HDBCN activity. **(A)** Representative image from a video recording synchronized with fiber-photometry measurements of VP and HDB cholinergic neuron activity. Pupil size was tracked using DeepLabCut, trained to identify pupil edges (P1–P3, P1’–P3’) and eyelid positions (L1–L1’). **(B)** Representative traces showing normalized pupil size (black) and normalized cholinergic activity in the VP (teal) and HDB (red). The boxed region is expanded in panel C. **(C)** Magnified view of the boxed region in panel B, illustrating that both VP and HDB cholinergic activity peaks are strongly synchronized with periods of pupil dilation. **(D)** Average pupil size triggered by transient calcium peaks of VPCNs and HDBCNs (n = 12 mice). **(E)** Population statistics comparing transfer entropy, which quantifies directional information flow between pupil size and calcium activity of VPCNs and HDBCNs. **(F)** Significance matrix for the transfer entropy analysis shown in panel E. Note that TE values from HDB or VP to pupil are not significantly different (p = 0.93, Mann-Whitney U-test).

We found that most pVPCNs showed precise reward and punishment responses similar to what was shown for BFCNs (Hangya et al., 2015; Laszlovszky et al., 2020; Hegedüs et al., 2023) (Fig.6B-D). Most pVPCNs were activated by rewards, punishments and conditioned stimuli, while a smaller population showed activation by positive valence and inhibition by negative valence stimuli (Fig.6C-G). These responses were unlike those described for CINs, especially regarding the well-characterized “pause-burst” reward responses of CINs in dorsal striatum (Inokawa et al., 2010; Zhang et al., 2018; Cox and Witten, 2019). Nevertheless, a few pVPCNs showed more delayed and sustained reward-elicited firing rate increase resembling those of CINs (Fig. S2), suggesting that while most VPCNs are BFCN-like, they co-occur with a smaller CIN-like population. These two types could even be recorded concurrently on the same electrode, suggesting that they are spatially mixed.

We also examined the autocorrelation of pVPCNs. Consistent with our in vitro recording, most pVPCNs showed a bursting in vivo firing pattern with a refractory period that was somewhat longer than what was previously shown for burst-BFCNs (Fig.6H-I) (Laszlovszky et al., 2020). Indeed, when we categorized pVPCNs to strongly bursting Burst-pVPCNs (Burst-pVPCN-SB), Poisson-like Burst-pVPCNs (Burst-pVPCN-PL) and regular rhythmic pVPCNs (Reg-pVPCNs) based on their burstiness and refractory period as was done for BFCNs (Laszlovszky et al., 2020), we found n = 10/21 Burst-pVPCN-SB and n = 10/21 Burst-pVPCN-PL neurons, a firing pattern distribution resembling HDBCNs (Fig.6J-K). Corroborating that a small fraction of VPCNs might be CIN-like, one pVPCN showed long refractory and theta-rhythmicity characteristic of both CINs and reg-BFCNs (n = 1/21 Reg-pVPCN).

### Pupil size correlates with both VPCN and HDBCN activity

Changes in pupil diameter under constant illumination were shown to be predicted by cholinergic transients originating from the basal forebrain (Nelson and Mooney, 2016; Reimer et al., 2016; Jing et al., 2020; Neyhart et al., 2024). We tested whether VPCNs resemble BFCNs in this aspect as well, by monitoring pupil diameter in parallel with VPCN and HDBCN bulk calcium signals (n = 12 mice, Fig.7A-C).

As expected, pupil dilations were temporally predicted by calcium transients recorded in HDBCNs of the basal forebrain. Similarly, we found that VPCN calcium peaks showed a comparable level of correlation with oncoming pupil dilations (Fig.7B-D). To perform a quantitative comparison of the predictive value of VPCN and HDBCN signals, we calculated transfer entropy (TE), an information theory measure of predictability across time series that is not restricted in the linear domain (Gourévitch and Eggermont, 2007) (Fig.7E). As expected based on the above temporal dynamics, prediction of the pupil size based on cholinergic signals (HDB to pupil TE and VP to pupil TE) were characterized by the highest TE values. At the same time, VPCNs showed comparable predictive values in terms of pupil size as the HDBCNs (p = 0.93, Mann-Whitney U-test), indicating that VPCNs are basal forebrain-like in their relation to pupil dilations.

## Discussion

By systematically comparing them to BFCNs, we demonstrated that most VPCNs belong to the basal forebrain cholinergic projection system based on their hodology, intrinsic biophysical properties and in vivo physiological responses to behaviorally salient appetitive and aversive events.

While not previously characterized, expectations about VPCN input-output connectivity are set by two classes of studies: tract tracing experiments of VP neurons and those of BFCNs. The mediodorsal thalamus is considered a primary output of VP, along with parts of the reticular and paraventricular thalamic regions (Zahm et al., 1996; Tripathi et al., 2013; Root et al., 2015). The VP also sends important projections to the lateral habenula and the VTA, which were shown to express PV and contain both GABAergic and glutamatergic components, linked to different aspects of depression (Knowland et al., 2017). Additionally, the VP sends topographically organized projections to the lateral hypothalamus and GABAergic efferents to the subthalamic nucleus, and projects back robustly to the nucleus accumbens, its major source of afferents (Root et al., 2015; Soares-Cunha et al., 2022; Domingues et al., 2023). While most of these projections are considered GABAergic, a strong cholinergic component of the VP to BLA pathway has been described (Root et al., 2015; Kim et al., 2024), while a cholinergic cortical projection was also assumed (Zaborszky et al., 2012). Concerning BFCNs, Do and colleagues characterized whole-brain distribution of axonal projections and found the hippocampus, piriform area, ventral striatum, amygdala and neocortical regions as main BFCN targets, though these were not stratified according to input cell location within the BF (Do et al., 2016). Except for an absence of hippocampal targets that are known to receive their cholinergic input from rostral BF (Agostinelli et al., 2019), we found VPCN projections consistent with BFCN outputs.

The densest input to VP is provided by GABAergic fibers from the nucleus accumbens, complemented by VTA/SNc dopaminergic, dorsal raphe serotonergic, STN glutamatergic, infralimbic cortical and amygdalar afferents (Root et al., 2015). It has been shown that besides dopaminergic, the VTA also provides GABAergic and glutamatergic VP inputs (Hnasko et al., 2012; Root et al., 2015). Whole-brain monosynaptic inputs to BFCNs were described by Hu et al. (Hu et al., 2016), pointing to the caudoputamen, central amygdala, lateral hypothalamus, nucleus accumbens and VTA as major sources of afferents, in line with earlier reports (Záborszky and Cullinan, 1992). We found that VPCNs received most of their monosynaptic inputs from nucleus accumbens, lateral hypothalamus and central amygdala, consistently with BFCNs in general. Thus, input-output mapping of VPCNs suggests that they are full-fledged members of the BF cholinergic projection system.

BFCNs were shown to be either early or late firing in acute slice experiments (Unal et al., 2012), and later demonstrated to form two distinct types of regular rhythmic and bursting neurons (Khateb et al., 1992; Alonso et al., 1996; Szymusiak et al., 2000) both in the nucleus basalis and in the HDB in vivo (Laszlovszky et al., 2020). Striatal cholinergic interneurons resemble Reg-BFCNs regarding their firing patterns in their slow-theta rhythmicity and long functional refractory period (Inokawa et al., 2010; Laszlovszky et al., 2020). We found that most VPCNs in vitro as well as pVPCNs in vivo showed bursting properties like Burst-BFCNs, with a few exceptions that fired like Reg-BFCNs and CINs. Of note, VPCNs showed slightly but distinctively longer refractory periods in their autocorrelaograms than BFCNs, the significance of which should be determined by future studies, These results again suggest that most VPCNs are BFCNs, in accordance with a topographical antero-posterior gradient of bursting cholinergic neurons within the BF (Laszlovszky et al., 2020).

VPCNs responded to rewards, punishments and reward-predicting stimuli, consistent with both BFCN (Lovett-Barron et al., 2014; Hangya et al., 2015; Harrison et al., 2016; Sturgill et al., 2020; Robert et al., 2021; Allard and Hussain Shuler, 2023; Hegedüs et al., 2023) and VP function in reward coding, motivation and associative learning (Tindell, 2004; Smith et al., 2009; Wassum et al., 2009; Richard et al., 2016b, 2018; Saga et al., 2017; Wulff et al., 2019; Ottenheimer et al., 2020a; Stephenson-Jones et al., 2020; Hegedüs et al., 2021; Soares-Cunha et al., 2022). When we performed a direct comparison with the HDB nucleus of the BF in Pavlovian conditioning, we found that VPCN and HDBCN calcium signals were robustly positively correlated. Nevertheless, bulk calcium recordings also revealed a bias in VPCNs toward reward responses, with faster and larger calcium responses to rewards but slower responses to punishments. This is in line with the known importance of VP in the reward aspects of learning (Tindell et al., 2006; Smith et al., 2009; Prasad et al., 2020), recent findings on the role of HDB in aversive coding and learning from negative experience (Hangya et al., 2015; Hegedüs et al., 2024), and supports the conclusion of Ottenheimer et al. that the VP processes certain aspects of reward independently of the nucleus accumbens, based on faster and more robust reward responses in the VP (Ottenheimer et al., 2018). Indeed, our results suggest that faster-than-striatal reward responses in the VP may arrive through VPCNs.

Spike responses of most pVPCNs to rewards, punishments and conditioned stimuli followed the known temporal dynamics of HDBCNs (Hangya et al., 2015; Sturgill et al., 2020; Hegedüs et al., 2023), while a few neurons exhibited striatal-like pause-burst responses to rewards (Inokawa et al., 2010). Most VPCNs showed correlated responses to positive and negative valence stimuli (Stephenson-Jones et al., 2020) similar to BFCNs (Hangya et al., 2015; Sturgill et al., 2020; Hegedüs et al., 2023), while a few VPCNs exhibited strong bias towards positive or negative outcomes, in line with a recent study that used olfactory stimuli (Kim et al., 2024). These results suggest an involvement of VPCNs in explicit learning likely including fear learning (Akmese et al., 2023; Ji et al., 2023), similar to what was shown for BFCNs in general (Letzkus et al., 2011; Jiang et al., 2016).

Changes in pupil size under constant illumination has been linked to multiple neuromodulatory systems (Larsen and Waters, 2018) including noradrenergic (Reimer et al., 2016; de Gee et al., 2017; Bang et al., 2023), cholinergic (Nelson and Mooney, 2016; Reimer et al., 2016; Jing et al., 2020; Neyhart et al., 2024) and serotonergic (Cazettes et al., 2021) activity, and were recently shown to reflect learning (Lee and Margolis, 2016) and consolidation processes (Chang et al., 2025) of associative memories. A difference in the time lag between acetylcholine rise and pupil dilation across different cortical areas suggested that different parts of the cholinergic system may have distinct temporal correlations with pupil size (Neyhart et al., 2024). We tested this by correlating VPCN and HDBCN activity with pupil diameter and found comparable predictive value of the two cholinergic signals in forecasting pupil dilations. This points to another functional aspect in which VPCNs act like BFCNs, further corroborating their functional similarity.

## Acknowledgements

We thank Katalin Lengyel for technical assistance in anatomical methods, and Anna Velencei and Jaishree Biswakarma for their contributions to animal training. We thank the FENS-Kavli Network of Excellence for fruitful discussions. This work was supported by the NKFIH K135561 and K147097 grants of the National Research, Development and Innovation Office, the Hungarian Brain Research Program NAP3.0 (NAP2022-I-1/2022) grant by the Hungarian Academy of Sciences, the “Lendület” LP2024-8/2024 grant by the Hungarian Academy of Sciences and the European Union project RRF-2.3.1-21-2022-00004 within the framework of the Artificial Intelligence National Laboratory.

---

**Supplementary figure 1.**
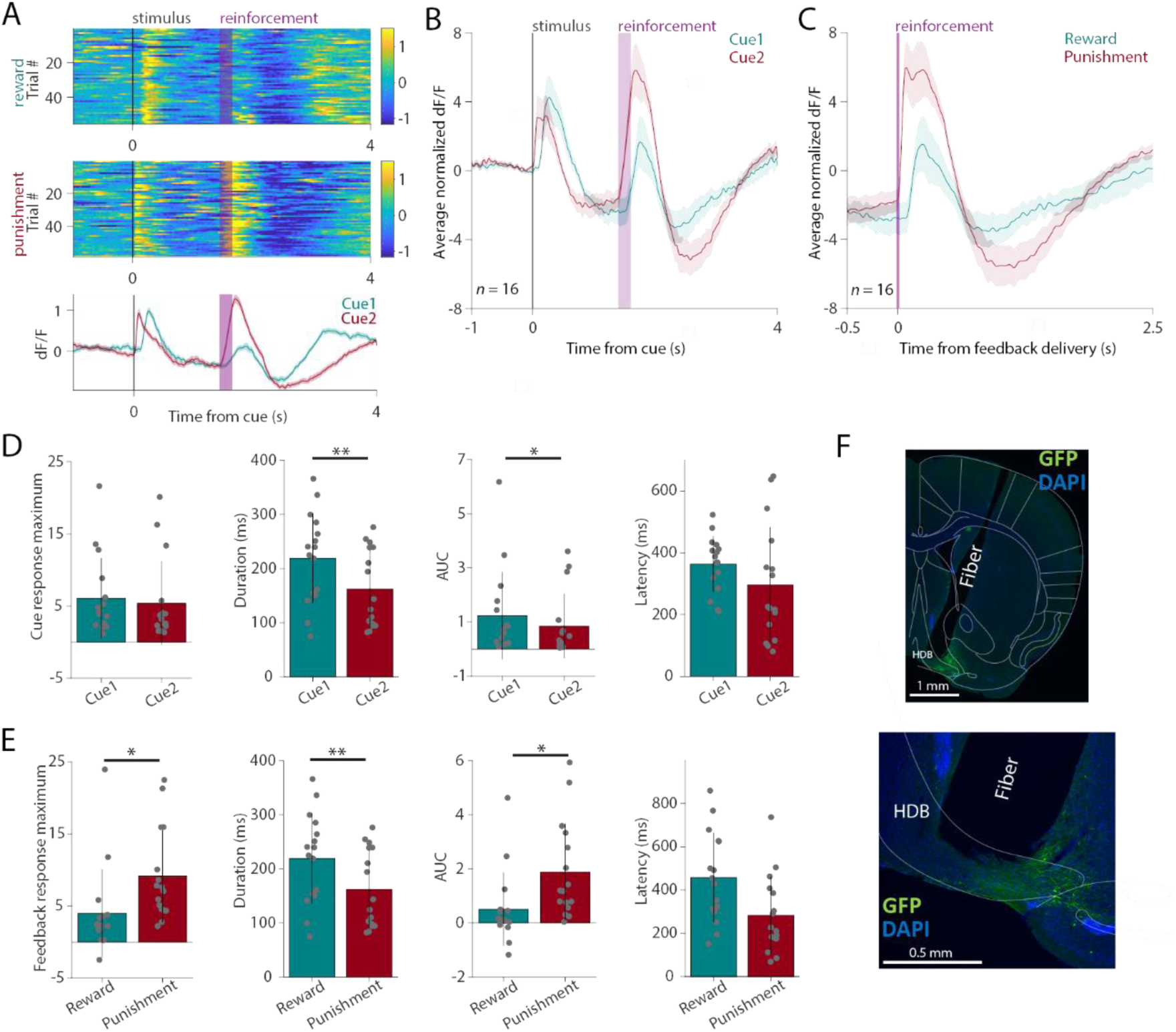
Cholinergic cells in the HDB respond differently to the reward- and punishment-predicting cues. **(A)** Example fiber photometry recording of HDBCNs from a single session. Top, normalized dF/F traces of all rewarded and punished trials aligned to cue onset, color coded (blue, low values; yellow, high values). Bottom, average dF/F traces from the same session. Error shades indicate SEM. **(B)** Average z-scored dF/F of HDBCNs aligned to the reward-predicting Cue 1 (green) and the punishment-predicting Cue 2 (red), averaged across all animals (n = 16). Error shades indicate SEM. **(C)** The same as in panel B but aligned to reward (green) and punishment delivery (red). Error shades indicate SEM. **(D)** From left to right, comparison of response magnitude, duration, integral and latency between HDBCN responses to the reward-predicting Cue 1and the punishment-predicting Cue 2. Each dot represents the session-average of a single animal. AUC, area under the curve. Bar graphs show mean ± standard deviation. *, p < 0.05; **, p < 0.01 Wilcoxon signed-rank test. **(E)** The same as in panel H but comparing HDBCN responses to reward and punishment. Each dot represents the session-average of a single animal. Bar graphs show mean ± standard deviation. *, p < 0.05; **, p < 0.01; Wilcoxon signed-rank test. **(F)** Representative fluorescent histological image of the measurement site. Scale bars, top, 1 mm; bottom, 0.5 mm.

**Supplementary figure 2.**
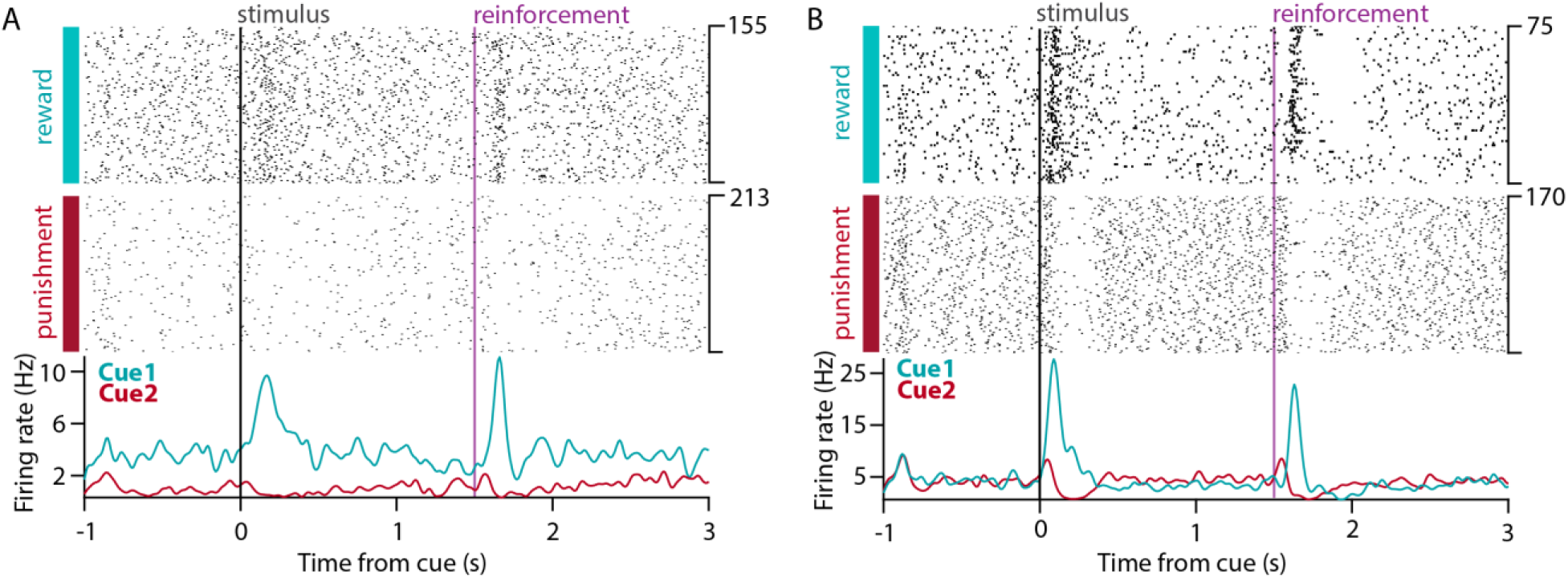
Few putative VPCNs show more delayed and sustained reward-elicited firing rate increase. **(A)** Top, raster plot of spike times aligned to cue onset of an example pVPCN that showed more delayed activation after the reward-predicting cue and the reward and no activation after the punishment-predicting cue and the punishment. Bottom, corresponding PETHs (green, rewarded trials; red, punished trials). **(B)** Same as in B, for a different example pVPCN.

**Table S1.** Abbreviations.

|  |  |
| --- | --- |
| AUC | area under the curve |
| ACG | autocorrelogram |
| AP | action potential |
| BFCNs | basal forebrain cholinergic neurons |
| BLA | basolateral amygdala |
| BNST | bed nucleus of the stria terminalis |
| Burst-BFCNs | bursting basal forebrain cholinergic neurons |
| Burst-pVPCN-SBs | strongly bursting putative ventral pallidal cholinergic neurons |
| Burst-pVPCN-PLs | Poisson-like bursting putative ventral pallidal cholinergic neurons |
| CCD | charge-coupled device |
| CCG | cross-correlogram |
| CeA | central amygdala |
| CINs | cholinergic interneurons |
| ChAT | choline acetyltransferase |
| DIC | differential interference contrast |
| eFEL | electrophys feature extraction library |
| HDB | horizontal limb of the diagonal band of Broca |
| HDBCNs | cholinergic neurons of horizontal limb of the diagonal band of Broca |
| HSA | human serum albumin |
| ISI | inter-spike interval |
| LH | lateral hypothalamus |
| LHb | lateral habenula |
| LS | lateral septum |
| MS | medial septum |
| NAc | nucleus accumbens |
| OD | outer diameter |
| PB | phosphate buffer |
| PETH | peri-event time histogram |
| PFA | paraformaldehyde |
| PFC | prefrontal cortex |
| POA | preoptic area |
| PSTN | parasubthalamic nucleus |
| pVPCNs | putative ventral pallidal cholinergic neurons |
| Reg-BFCNs | regular rhythmic basal forebrain cholinergic neurons |
| SE | standard error |
| SI | substantia innominata |
| SPL | sound pressure level |
| STN | subthalamic nucleus |
| TBS | tris buffered saline |
| TE | transfer entropy |
| TTL | transistor-transistor logic |
| UMAP | Uniform Manifold Approximation |
| VP | ventral pallidum |
| VPCNs | ventral pallidal cholinergic neurons |
| pVPCNs | putative ventral pallidal cholinergic neurons |
| VTA | ventral tegmental area |

**Table S2.** Primary antibodies used in immunhistochemical experiments.

| Antigen | Host | Dilution | Source | Catalog number | Specificity | Characterized in |
| --- | --- | --- | --- | --- | --- | --- |
| eGFP | Chicken | 1:2000 | Thermo Fisher Scientific | A10262 | No staining in mice not injected with eGFP-expressing virus. | Information of the distributor. |
| mCherry | Rabbit | 1:1000 | BioVision | 5993-100 | No staining in mice not injected with mCherry-expressing virus. | Information of the distributor. |
| ChAT | rabbit | 1:500 | Synaptic Systems | 297013 | Specific for rat and mouse ChAT. | Information of the distributor. |

**Table S3.** Secondary antibodies used in immunohistochemical experiments.

| Raised in | Raised against | Conjugated with | Dilution | Source | Catalog number |
| --- | --- | --- | --- | --- | --- |
| Donkey | Chicken | Alexa 488 | 1:1000 | Jackson ImmunoResearch | 703-545-155 |
| Donkey | Rabbit | Alexa 594 | 1:500 | Thermo Fisher Scientific | A21207 |

## Notes

### Competing Interest Statement

The authors have declared no competing interest.

